# Cerebellum-inspired Kernel for Efficient Out-of-Distribution Detection

**DOI:** 10.64898/2026.03.11.710983

**Authors:** Yaoduo Zhang, Jian Zhang, Yunliang Zang

## Abstract

Detecting novel stimuli is a fundamental neural function, yet its machine learning counterpart—out-of-distribution (OOD) detection—remains challenging, with models often making overconfident predictions on unseen inputs. Inspired by the strong pattern-separation capabilities of cerebellum-like circuits, we introduce a cerebellum-inspired kernel with an efficient closed-form implementation. Combining random Gaussian projection with Top-k sparsification, the kernel reshapes similarities in high-dimensional space to enhance separability between in-distribution (ID) and OOD samples. On OpenOOD benchmarks, our kernel consistently improves multiple baseline methods, and pairing it with the energy score achieves performance comparable to or exceeding current state-of-the-art approaches. The closed-form design also avoids the high computational cost of large-expansion explicit mapping. These results demonstrate the generality and potential of cerebellar kernels for OOD detection and other tasks requiring efficient pattern separation under limited computational resources.

## 1 Introduction

Artificial neural networks (ANN) have achieved remarkable progress in tasks such as image recognition. Models can perform well on unseen in-distribution (ID) samples; however, in real deployments, this assumption is frequently violated. For samples completely unrelated to the training data, the model still outputs a prediction, often making overconfident yet incorrect predictions when meeting out-of-distribution (OOD) samples. The misclassification can cause severe safety and risk issues. Therefore, OOD detection aims to distinguish ID samples from OOD samples. A common practice is to construct a scalar OOD score for each input and apply a threshold for rejection or alarms, improving the usability and trustworthiness of models in open-world settings [1].

Biological organisms—including humans—can rapidly and accurately recognize when an object or scene is novel, or when the environment has changed significantly. Although the biological mechanisms underlying this capability remain incompletely understood, the brain must contain structures that support functions analogous to OOD detection. Studies have suggested that cerebellum-like circuits—such as the cerebellum itself and the fly olfactory circuit—play a role in anomaly detection [2–5].

Classical cerebellar theory proposes that such circuits are well-suited for pattern separation, implying that the cerebellum’s feedforward architecture may be key to enhancing OOD detection in ANN. Algorithmically, a cerebellum-like feedforward structure can be abstracted as mapping low-dimensional features into a high-dimensional space via random Gaussian projection followed by a winner-take-all (Top-k) selection. This expansion can facilitate novelty recognition by making OOD detection easier.

However, pure pattern separation alone does not sufficiently explain the role of cerebellar structure in OOD detection. The reason is that similarity reduction occurs globally across all samples, whereas distinguishing OOD from ID requires differential changes in similarities among ID samples (within and between classes) and between ID and OOD samples. Therefore, a systematic theory is needed to explain how cerebellum-style expansion affects pairwise similarities.

In ANNs, one efficient OOD detection strategy is post-processing—using a trained classifier at test time to identify OOD samples based on logits, penultimate-layer features, or gradients [6, 7]. Many of these methods rely on inner products or cosine similarities. We propose mapping extracted features into a “cerebellar space” and performing similarity computations, thereby incorporating cerebellar principles into post-processing OOD detection.

A key challenge in directly embedding features into existing methods is that stable high-dimensional representations require a large expansion factor—often at least 20× the original feature dimension—resulting in substantial computational overhead [8]. To address this, we derive a cerebellum-inspired inner-product kernel, Ψ_ceb_(*x*), that computes the inner product between mapped features in cerebellar space directly from sufficient statistics of the original feature pair, without explicitly performing the mapping.

We define the explicit cerebellum-style expansion mapping as

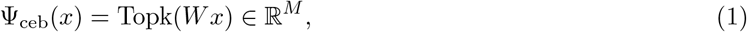

where *W* ∈ ℝ^*M* ×*N*^ is a random projection matrix with i.i.d. entries *W*_*ji*_ ∼ 𝒩 (0, 1), *M* is the expanded dimension, *N* is the input dimension and Topk(·) keeps the *k* largest elements with *k* determined by the retention ratio *s* (Top-k ratio), where *k* = *sM*. Biologically, cerebellum-like circuits use sparse connectivity; for a stable and simple engineering implementation we use dense Gaussian projection, under which the probabilistic analysis in Section 3 is exact. This explicit mapping induces kernels through inner products or cosine similarities in the mapped space.

Our closed-form formulation significantly improves efficiency, reducing complexity from *O*(*MN*) to *O*(*N*). The kernel corresponds to the infinite-expansion limit of the explicit mapping, so numerical values may differ slightly from finite computations, but the conceptual equivalence remains. The cerebellar kernel itself introduces only one specific hyperparameter, the Top-k ratio, allowing for concise theoretical analysis.

To evaluate whether cerebellum-style feature expansion and the proposed kernel can improve OOD detection at the feature level, we integrate it into multiple OOD baselines. We first introduce CK-Energy, a method based on the energy score (EBO) [9] that implicitly leverages cerebellar high-dimensional features, and then examine its combination with other OOD detection approaches. Across diverse datasets, we observe consistent performance gains, demonstrating that the cerebellar kernel improves ID/OOD separability and is broadly applicable.

Our main contributions are as follows:

1. **Closed-form cerebellar kernel:** Starting from a feature mapping inspired by cerebellar feedforward expansion, we derive closed-form functions for computing inner products and cosine similarities in cerebellar space directly from the original features. We further optimize the computation to achieve *O*(*N*) complexity, eliminating the overhead of explicit high-dimensional mapping and ensuring high efficiency for practical OOD detection.
2. **Broad applicability:** The proposed cerebellar kernel can be integrated with various OOD detection methods—including, but not limited to, similarity-based approaches—to improve their performance. Combining the energy score with the cerebellar kernel yields performance and robustness competitive to or exceeding current SOTA methods.
3. **Analytical insights:** The kernel formulation enables clear analysis of the feature mapping Ψ_ceb_(*x*) and the induced kernel space, providing theoretical understandings of why cerebellum-inspired structures benefit OOD detection.

## 2 Related Work

### Cerebellar theory

Classical theories of cerebellar computation view the cerebellar cortex as implementing a massive expansion recoding that supports robust discrimination and learning from error signals [2, 3, 10]. Sparse, high-dimensional expansions combined with competitive selection (winner-take-all) have been argued to improve separability and reduce interference [11, 12], and related “expansion + sparsification” motifs also appear in other biological circuits [4]. These ideas connect naturally to machine learning notions of sparse distributed representations and associative memory [13–15].

### Out-of-distribution detection

OOD detection is often approached via post-hoc scoring of trained classifiers, with early baselines using maximum softmax probability [1]. Logit calibration and input preprocessing, as in ODIN [6], improve separation, while gradient-based analysis, e.g., GradNorm [16], utilizes the magnitude of gradients to distinguish OOD samples. Feature-space statistics, such as class-conditional Gaussian modeling with Mahalanobis distance [7], and energy-based scores [9] unify confidence measures. Representation-shaping methods like ReAct [17], ASH [18], and Gram-matrix features [19] highlight the role of intermediate feature geometry. Subspace-based approaches, e.g., ViM [20], connect to classical kernel methods [21]. Beyond post-hoc scoring, Outlier Exposure [22] improves robustness when auxiliary outliers are available. Uncertainty estimation via Bayesian approximations [23] and ensembles [24] remains standard, though calibration studies reveal persistent overconfidence under shift [25, 26]. Recent benchmarks, e.g., OpenOOD and ImageNet OOD audits, have standardized protocols and exposed evaluation pitfalls [27, 28].

## 3 Preliminaries

We summarize the probabilistic setup and notation used to derive the closed-form cerebellar kernel.

### 3.1 Cerebellar expansion and quantile gating

Following Eq. (1), we consider an explicit expansion matrix *W* with i.i.d. Gaussian entries. Denote the *j*-th row by 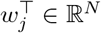, where *w*_*j*_ ∼ 𝒩(0, *I*_*N*_). For an input *x* ∈ ℝ^*N*^, the explicit expansion computes row responses 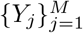,

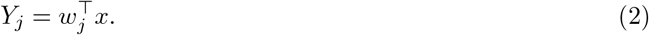

Top-k gating then keeps the *k* = *sM* largest responses among 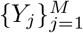 (with retention ratio *s* ∈ (0, 1)). For analysis, we approximate Top-k by a fixed quantile threshold on standardized responses. Since *Y*_*j*_ | *x* ∼ 𝒩(0, ∥*x*∥^2^), letting *Z* = *Y*/∥*x*∥ ∼ 𝒩(0, 1) and setting

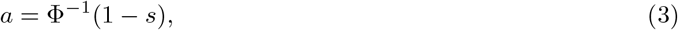

we define the single-row gated output

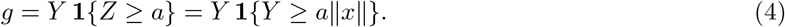

This approximation becomes exact as *M* → ∞; details are referred to Appendix A.1. With *g*_*j*_(*x*) defined per row, we use the globally normalized implicit feature

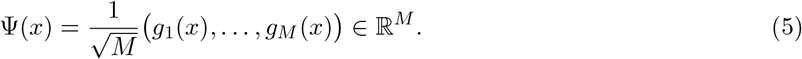

Since rows are i.i.d. and we use global 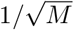 normalization, for sufficiently large *M*, the explicit inner product concentrates around the single-row expectation 𝔼[*gg*′].

### 3.2 Gaussian projection and truncated moments

For two inputs *x, x*′ ∈ ℝ^*N*^, define the basic statistics

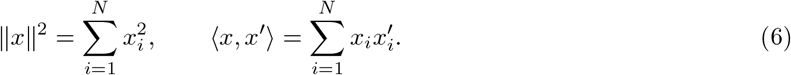

Under the Gaussian projection *Y* = *w*^⊤^*x* and *Y* ′ = *w*^⊤^*x*′ with *w* ∼ 𝒩(0, *I*_*N*_), we have

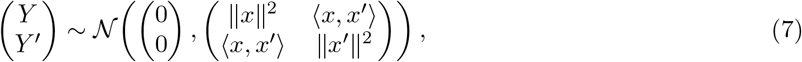

and therefore the standardized variables *Z* = *Y*/ ∥*x*∥ and *Z*′ = *Y* ′ / ∥*x*′∥ satisfy (*Z, Z*′) ∼ 𝒩 (0, Σ_*ρ*_) with correlation

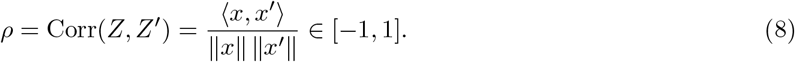

Let *I* = **1** {*Z* ≥*a*} (and *I*′ analogously), where *a* is determined by the retention ratio *s* in Eq. (3). We use the one-dimensional truncated moments

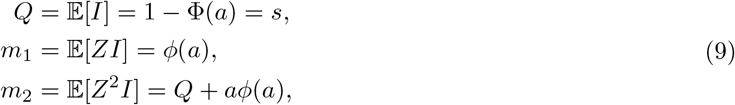

and the two-dimensional truncated moments for (*Z, Z*′) ∼ 𝒩(0, Σ_*ρ*_)

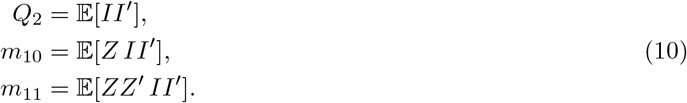

Their integral forms and numerical evaluation are provided in Appendix A.2.

## 4 Method

We propose a closed-form kernel to retain the expressivity of cerebellar mapping while eliminating the overhead of explicit high-dimensional feature generation. Unlike explicit methods where complexity scales with dimension *M* despite performance saturation [8], our kernel computes the infinite-expansion limit directly. This formulation decouples computational complexity from *M*. We derive these limiting kernels (Section 4.1), detail their efficient implementation (Section 4.2), and integrate them with OOD baselines (Section 4.3).

### 4.1 Closed-form cerebellar kernel

Many post-processing OOD detection methods use penultimate-layer features extracted from a classifier to compute OOD scores, often involving inner products ⟨*x, x*′⟩ or cosine similarity 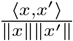. We seek to apply the cerebellum-style mapping Ψ_ceb_(*x*) in this stage. Under the setup in Section 3 (Gaussian projection and quantile-threshold approximation), both the inner product and cosine similarity of the cerebellar kernel can be written using truncated bivariate normal moments, enabling an efficient closed-form implementation.

In our Gaussian projection setting, *Y* = *w*^⊤^*x* and *Y* ′ = *w*^⊤^*x*′ are jointly normal with zero mean, so the induced kernel depends on the original pair only through its cosine correlation 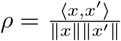 (Section 3). Let (*Z, Z′*) = (*Y*/∥*x*∥, *Y*′/∥*x*′ ∥) ∼ 𝒩(0, Σ_*ρ*_), and for Pos-Topk (Top-k of positive) use the standardized threshold *a* = Φ^−1^(1 − *s*) with *I* = **1** {*Z* ≥ *a*} and *I*′ = **1** {*Z*′ ≥ *a*}. Using the truncated moments *m*_2_(*a*) = 𝔼[*Z*^2^*I*] = *Q* + *aϕ*(*a*) and *m*_11_(*a, ρ*) = 𝔼[*ZZ*′*II*′] defined in Section 3, the infinite-expansion Pos-Topk kernels become

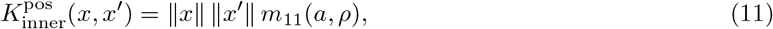

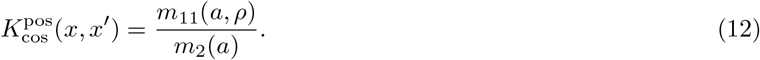

#### Cerebellar kernel with Abs-Topk

One-sided Top-k (Pos-Topk) discards large negative activations, which may also carry discriminative information (e.g., inhibitory responses). Since the essence of Topk is to retain the most salient activations as representatives, we introduce absolute-value Top-k (Abs-Topk): select Top-k by element-wise magnitude |*x*|, thus preserving representative information on both positive and negative sides. When computing inner products, we keep the original signs, improving symmetry and expressiveness.

We adopt gating on standardized magnitude |*Z*|, whose threshold depends only on the retention rate *s* and is convenient for batch computation and tabulation (the alternative |*Y*| gating is discussed in Appendix A.3). Let

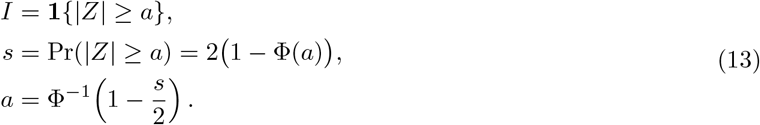

By symmetry, 𝔼[*ZI*] = 0. Let 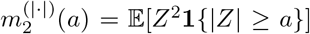 and 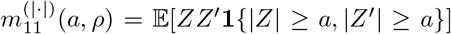 denote the corresponding truncated moments (Appendix A.3). The Abs-Topk kernels are

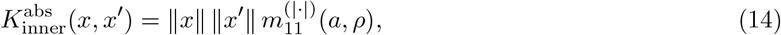

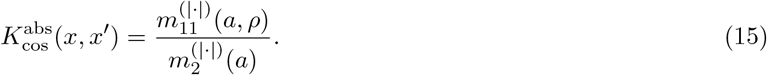

According to these kernels, the correlation *ρ* of the original feature pair is the most critical factor affecting inner products and cosine similarities in cerebellar space. Through the truncated-moment integrals, *ρ* adjusts the coefficients of the sufficient statistics and thus the final output.

Based on the closed-form expression, the impact of cerebellar kernels on similarity depends on input correlations. As shown in Figure 1 (Abs-Topk as an example), pairs with lower original correlations (e.g., between classes) experience a larger relative reduction after applying the cerebellar kernel. This differential similarity reduction enhances ID-OOD separation in kernel-space geometry, easing OOD detection.

**Figure 1:**
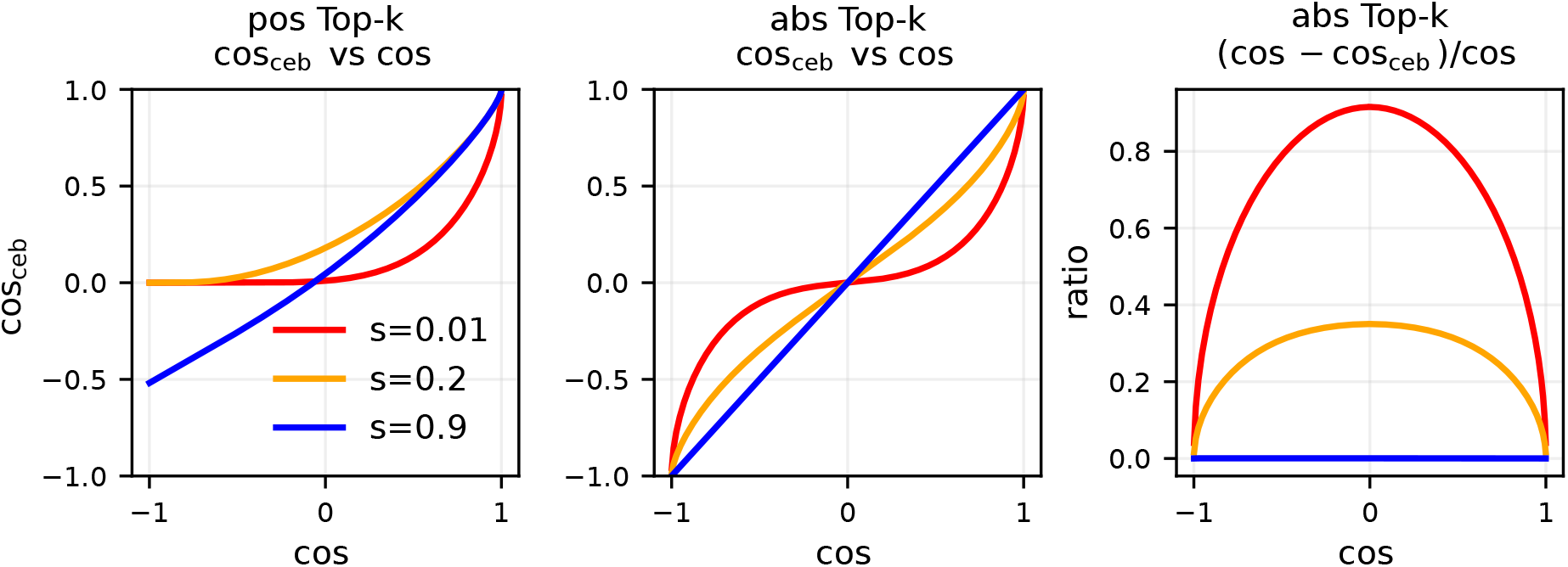
Visualization of cosine output vs. pairwise correlation (left, Pos-Topk; middle, Abs-Topk) and the relative correlation reduction for Abs-Topk kernels.

### 4.2 Efficient computation and engineering implementation

The closed-form expression avoids explicitly generating *M* -dimensional features and the expensive row-wise Top-k operation. The remaining challenge is that the key term *m*_11_(*a, ρ*) is defined by a one-dimensional integral (Appendix A.2), and evaluating it online for every pair would dominate the cost. For a fixed retention rate *s*, the threshold *a* is fixed, and *m*_11_(*a, ρ*) depends only on *ρ*. We therefore precompute this function offline and use *O*(1) interpolation online, yielding a two-stage pipeline: offline tabulation of truncated moments + online interpolation lookup.

For one-sided Top-k gating (*Z* ≥ *a*), given *s* we have *a* = Φ^−1^(1 − *s*). The one-dimensional integral expression for *m*_11_(*a, ρ*) is given in Appendix A.2. On a uniform grid 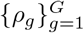 over *ρ* ∈ [−*ρ*_max_, *ρ*_max_] (typically choosing *ρ*_max_ *<* 1 to avoid numerical instability in 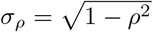), we apply Simpson integration and store

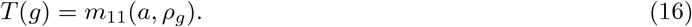

At test time, for any *ρ*, we locate the neighboring grid points and perform linear interpolation to obtain 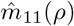. Abs-Topk(|*Z*|) reuses the same table at *ρ* and −*ρ* and combines them using the identities in Appendix A.3, requiring no extra integration.

To compute the cerebellar kernel for an input pair (*x, x*′), we compute ∥*x*∥, ∥*x*′∥ and 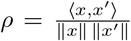 in *O*(*N*) time, query 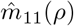 via interpolation, and combine it with the closed-form normalization terms (*m*_2_ for Pos-Topk, and 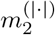 for Abs-Topk). The pseudocode is given in Algorithm 1.

#### Algorithm 1 Closed-form computation of the cerebellar kernel (inner product / cosine).

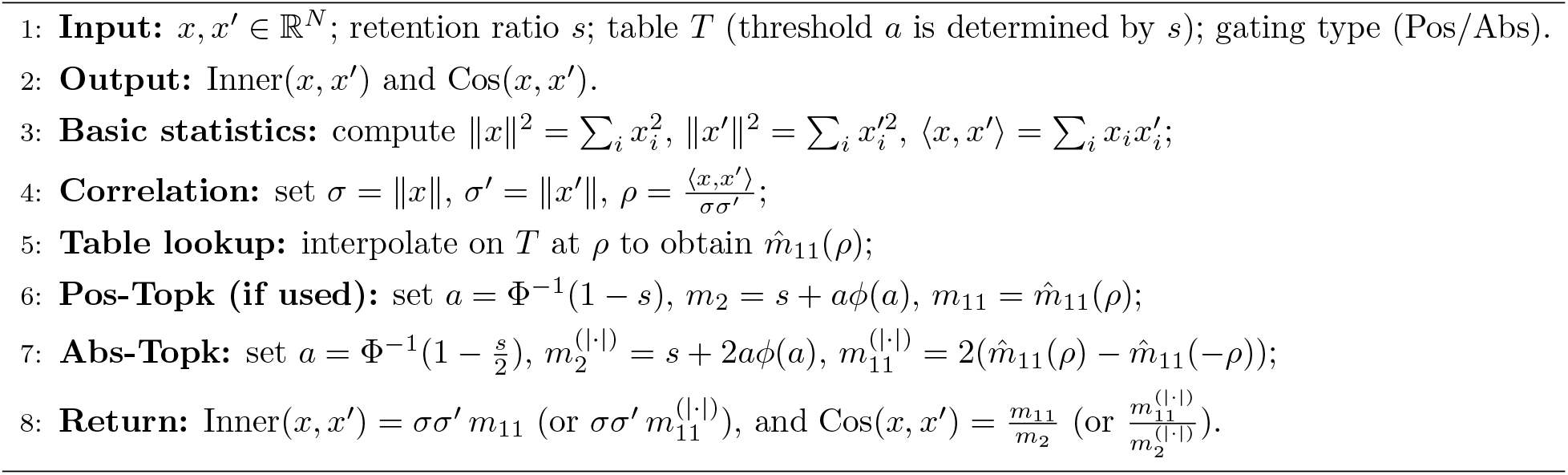

In the online stage, the dominant cost per input pair is *O*(*N*) (computing ∥*x*∥, ∥*x*′∥, ⟨*x, x*′⟩), while interpolation and coefficient combination are *O*(1). Offline tabulation costs *O*(*G* · grid), where grid is the number of quadrature points used per one-dimensional integral (per grid point *ρ*_*g*_).

### 4.3 Applying the cerebellar kernel to OOD detection

To examine the role of the cerebellar kernel in OOD detection, we integrate it with different methods to assess its performance and generality. We construct three models based on (i) energy scores (EBO), (ii) subspaces (PCA), and (iii) memory retrieval (HE). Our main model, CK-Energy (CKE), builds on EBO and is described in detail for subsequent analysis. The other two models, KPCA and KHE, extend PCA and HE, respectively, and are detailed in the Appendix, with comparisons to their baselines provided in Section 5.3.

In EBO, energy is computed from logits produced by a forward pass. To leverage the properties of cerebellar high-dimensional features, we need new features to produce new logits. We build a kernel classification head: we freeze all parameters except the last layer and replace the last layer with a kernel head. For each class *c*, we learn parameters (*w*_*c*_, *b*_*c*_) and compute a new classification logit via the Abs-Topk cosine kernel:

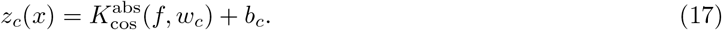

Here *f* denotes the feature extracted by the backbone classifier. Using all training features and labels, we minimize cross-entropy (CE) ℒ = CE(softmax(*z*(*x*)), *y*). We use the cerebellar kernel with cosine similarity, while we also observe that the inner-product variant achieves comparable performance. We then compute the energy from the new logits:

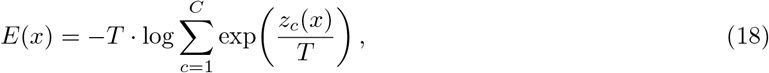

and set the OOD score as *S*(*x*) = −*E*(*x*) to match the benchmark convention. In this way, we implicitly leverage the geometry of the cerebellar kernel space.

For the subspace-based method, we use the reconstruction error of a test sample on the principal subspace of the ID training set as the OOD score, reformulated as reconstruction error in kernel space. We also build class-specific submodels and match each test sample to the submodel corresponding to its predicted class to reduce the computational burden of KPCA. For the memory-retrieval method, we follow HE [29] and replace the inner-product computation with 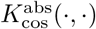, which is equivalent to retrieval and matching in the mapped feature space. Details are provided in the Appendix.

## 5 Experiments

### 5.1 Experimental Setup

To validate the effectiveness of our method, we follow the standard OpenOOD protocol and data splits [27], and report results on three common ID benchmarks: ImageNet-200, ImageNet-1k [30], and CIFAR-100 [31]. We evaluate our method under two distinct settings: **Standard OOD** and **Full-Spectrum OOD (FSOOD)**.

#### Standard OOD Setting

Following OpenOOD, dataset groups are divided into Near-OOD and Far-OOD. When ImageNet-200 or ImageNet-1k is used as the ID dataset, Near-OOD includes SSB-hard and NINCO; Far-OOD includes iNaturalist, Textures, and OpenImage-O [28, 32–34]. When CIFAR-100 is used as the ID dataset, Near-OOD includes CIFAR-10 and TinyImageNet (TIN); Far-OOD includes MNIST, SVHN, Textures, and Places365 [33, 35–38].

#### FSOOD Setting

The FSOOD benchmarks are designed to test OOD detection under a wide range of distribution shifts. In this setting, the OOD datasets remain consistent with the Standard OOD benchmark, while the ID set is adjusted by augmenting it with covariate-shifted samples—ImageNet-C [39], ImageNet-R [40], and ImageNet-V2 [41]—which are treated as ID together with clean validation data.

#### Implementation and Metrics

In standard OOD and FSOOD, we use the official model architectures and pretrained weights. For CIFAR-100 (not in FSOOD) and ImageNet-200, we use ResNet-18; for ImageNet-1k, we use ResNet-50 [42]. We report two widely used OOD metrics: AUROC (higher is better) and FPR@95 (lower is better), measuring overall separability and the false-positive rate at 95% true-positive rate, respectively. For each metric, we report the average performance on Near-OOD and Far-OOD as well as the overall average (ALL).

### 5.2 OOD detection performance in CKE

We systematically compare CKE with a range of classical post-processing methods under Near-OOD, Far-OOD, and ALL settings. Table 1 summarizes the standard OOD results on the ImageNet benchmarks, with CIFAR-100 results provided in the Appendix. Table 2 presents the FSOOD results on the ImageNet benchmarks.

**Table 1:** Results on the **Standard OOD** ImageNet benchmarks (Near/Far/ALL summary).

| Method | ImageNet-200 |  |  |  |  |  | ImageNet-1k |  |  |  |  |  |
| --- | --- | --- | --- | --- | --- | --- | --- | --- | --- | --- | --- | --- |
|  | Near-OOD |  | Far-OOD |  | ALL |  | Near-OOD |  | Far-OOD |  | ALL |  |
|  | FPR@95 | AUROC | FPR@95 | AUROC | FPR@95 | AUROC | FPR@95 | AUROC | FPR@95 | AUROC | FPR@95 | AUROC |
| OpenMax | 63.48 | 80.27 | 33.12 | 90.20 | 45.26 | 86.23 | 69.07 | 74.77 | 34.31 | 89.26 | 48.21 | 83.46 |
| ODIN | 66.76 | 80.27 | 34.23 | 91.71 | 47.24 | 87.13 | 72.50 | 74.75 | 43.96 | 89.47 | 55.38 | 83.58 |
| MDS | 79.11 | 61.93 | 61.66 | 74.72 | 68.64 | 69.60 | 85.45 | 55.44 | 62.92 | 74.25 | 71.93 | 66.73 |
| MDSEns | 91.75 | 54.32 | 80.96 | 69.27 | 85.28 | 63.29 | 93.52 | 49.67 | 82.77 | 67.52 | 87.07 | 60.38 |
| RMDS | **54.02** | 82.57 | 32.45 | 88.06 | 41.08 | 85.86 | 65.04 | 76.99 | 40.91 | 86.38 | 50.56 | 82.62 |
| Gram | 86.40 | 67.67 | 84.36 | 71.19 | 85.18 | 69.78 | 86.63 | 61.70 | 67.36 | 79.71 | 75.07 | 72.51 |
| EBO | 60.24 | 82.50 | 34.86 | 90.86 | 45.01 | 87.52 | 68.56 | 75.89 | 38.39 | 89.47 | 50.46 | 84.04 |
| ReAct | 62.49 | 81.87 | 28.50 | 92.31 | 42.10 | 88.13 | 66.69 | 77.38 | 26.31 | 93.67 | 42.46 | 87.15 |
| VIM | 59.19 | 78.68 | 27.20 | 91.26 | 40.00 | 86.23 | 71.35 | 72.08 | 24.67 | 92.68 | 43.34 | 84.44 |
| KNN | 60.18 | 81.57 | 27.27 | 93.16 | 40.43 | 88.52 | 70.87 | 71.10 | 34.13 | 90.18 | 48.83 | 82.55 |
| ASH | 64.89 | 82.38 | 27.29 | 93.90 | 42.33 | 89.29 | 63.32 | 78.17 | 19.49 | 95.74 | 37.02 | 88.71 |
| SHE | 66.80 | 80.18 | 42.17 | 89.81 | 52.02 | 85.96 | 73.01 | 73.78 | 41.45 | 90.92 | 54.07 | 84.06 |
| GEN | 55.20 | **83.68** | 32.10 | 89.81 | 41.34 | 87.36 | 65.32 | 76.85 | 35.61 | 89.76 | 47.49 | 84.60 |
| CKE | 55.68 | 83.23 | **20.75** | **94.96** | **34.70** | **90.27** | **62.59** | **80.43** | **14.59** | **96.63** | **33.79** | **90.15** |

**Table 2:** Results on the **FSOOD** ImageNet benchmarks (Near/Far/ALL summary).

| Method | ImageNet-200 |  |  |  |  |  | ImageNet-1k |  |  |  |  |  |
| --- | --- | --- | --- | --- | --- | --- | --- | --- | --- | --- | --- | --- |
|  | Near-OOD |  | Far-OOD |  | ALL |  | Near-OOD |  | Far-OOD |  | ALL |  |
|  | FPR@95 | AUROC | FPR@95 | AUROC | FPR@95 | AUROC | FPR@95 | AUROC | FPR@95 | AUROC | FPR@95 | AUROC |
| OpenMax | 88.27 | 50.89 | 74.10 | 67.96 | 79.77 | 61.13 | 81.59 | 57.03 | 57.44 | 75.24 | 67.10 | 67.96 |
| ODIN | 89.18 | 48.33 | 73.55 | 66.25 | 79.81 | 59.09 | 83.21 | 57.41 | 63.29 | 75.62 | 71.26 | 68.34 |
| MDS | 91.21 | 52.62 | 77.41 | 68.00 | 82.93 | 61.85 | 91.15 | 46.02 | 73.83 | 67.02 | 80.76 | 58.62 |
| MDSEns | 96.06 | 37.90 | 89.91 | 53.18 | 92.37 | 47.07 | 94.89 | 42.47 | 86.25 | 59.98 | 89.71 | 52.98 |
| RMDS | **84.33** | 58.59 | 72.39 | 66.62 | 77.17 | 63.41 | 78.69 | 62.06 | 61.61 | 73.29 | 68.44 | 68.79 |
| Gram | 86.98 | **61.23** | 85.41 | 65.26 | 86.04 | 63.65 | 88.55 | 56.28 | 72.13 | 75.20 | 78.70 | 67.63 |
| EBO | 87.16 | 50.51 | 75.71 | 64.59 | 80.29 | 58.96 | 81.17 | 56.61 | 60.59 | 73.37 | 68.82 | 66.67 |
| ReAct | 87.91 | 50.55 | 71.72 | 67.73 | 78.20 | 60.86 | 79.94 | 59.93 | 50.00 | 82.78 | 61.97 | 73.64 |
| VIM | 86.98 | 51.22 | 67.06 | 73.19 | 75.03 | 64.40 | 83.35 | 52.50 | 48.82 | 80.10 | 62.63 | 69.06 |
| KNN | 86.48 | 49.28 | 70.86 | 71.84 | 77.11 | 62.82 | 82.64 | 51.82 | 56.69 | 76.02 | 67.07 | 66.34 |
| ASH | 87.25 | 54.74 | 66.95 | 75.48 | 75.07 | 67.18 | **77.61** | 60.52 | 42.50 | 86.75 | 56.54 | 76.25 |
| SHE | 88.82 | 54.73 | 76.65 | 70.47 | 81.52 | 64.18 | 82.78 | 61.21 | 57.50 | 83.04 | 67.62 | 74.31 |
| GEN | 84.85 | 51.59 | 73.24 | 65.46 | 77.89 | 59.91 | 79.05 | 57.84 | 58.23 | 74.31 | 66.56 | 67.72 |
| NAC | 90.79 | 49.61 | 73.73 | 67.01 | 80.56 | 60.05 | 83.03 | 59.48 | 41.52 | 87.48 | 58.12 | 76.28 |
| KAN | 91.01 | 59.74 | 70.83 | 79.28 | 78.90 | 71.46 | 82.11 | 62.71 | 49.37 | 89.05 | 62.47 | 78.52 |
| CKE | 85.57 | 57.83 | **61.21** | **82.85** | **70.95** | **72.84** | 77.72 | **66.04** | **31.59** | **91.64** | **50.04** | **81.40** |

Overall, CKE maintains a consistent advantage over competing approaches across all benchmarks: it consistently reduces false positives (FPR@95) and improves separability (AUROC) on Far-OOD, while preserving competitive performance on Near-OOD, yielding the highest overall average performance.

In the standard OOD setting, CKE demonstrates superior efficacy on ImageNet-200, achieving the highest Far-OOD performance with an AUROC of 94.96% and an FPR@95 of 20.75%. This substantial gain over other baselines does not compromise Near-OOD detection, where CKE remains highly competitive against other top-performing methods, resulting in the highest overall average performance (AUROC = 90.27%, FPR@95 = 34.70%).

The advantage of CKE is even more pronounced on the larger-scale ImageNet-1k benchmark, dominating in Far-OOD (AUROC = 96.63%, FPR@95 = 14.59%), Near-OOD (AUROC = 80.43%, FPR@95 = 62.59%), and overall average performance (AUROC = 90.15%, FPR@95 = 33.79%). On CIFAR-100, CKE also delivers balanced and effective detection across both categories. Detailed breakdowns are provided in Tables 7, 8, and 11 of the Appendix.

We further evaluate model robustness under covariate shifts and distribution changes using the FSOOD benchmarks. CKE is compared against the strongest baselines in this setting—specifically KAN and ASH, which represent the previous state-of-the-art for robust OOD detection. CKE demonstrates remarkable resilience, achieving the best Far-OOD and overall average performance on both ImageNet-200 and ImageNet-1k.

On ImageNet-200, CKE outperforms other methods in both metrics. In Far-OOD, it achieves an AUROC of 82.85% and FPR@95 of 61.21%, compared to KAN’s 79.28% and 70.83% and ASH’s 75.48% and 66.95%. For the overall average, CKE reaches an AUROC of 72.84% and FPR@95 of 70.95%, surpassing KAN’s 71.46% and 78.90% and ASH’s 67.18% and 75.07%. A similar trend is observed on ImageNet-1k, where CKE consistently outperforms KAN and ASH in both Far-OOD and overall average performance. These results confirm that the cerebellar kernel enhances robustness against substantial distribution shifts. Full per-dataset results are provided in Tables 9 and 10 of the Appendix.

### 5.3 Combining cerebellar kernels with multiple methods

We hypothesize that enhancing ID/OOD separability via the cerebellar kernel can benefit a broad range of models. Besides EBO, we evaluate its generality by applying the cerebellar kernel to PCA and HE on the CIFAR-100 benchmark (Table 3).Results show that the cerebellar kernel improves the performance of all three methods, with the largest gain for the subspace-based PCA approach: AUROC increases by 6.89% and FPR@95 decreases by 13.22% in overall average performance. This suggests that the cerebellar kernel alters the subspace structure of ID/OOD data, making their subspaces more distinct to facilitate OOD detection. The other two methods also exhibit significant gains.

**Table 3:** Comparison between multiple baselines and their cerebellar-kernel variants on CIFAR-100.

| Method | Near-OOD |  | Far-OOD |  | ALL |  |
| --- | --- | --- | --- | --- | --- | --- |
|  | FPR@95 | AUROC | FPR@95 | AUROC | FPR@95 | AUROC |
| EBO | 59.21 | **80.91** | 56.59 | 79.77 | 56.26 | 80.15 |
| CKE | **55.43** | 78.76 | **46.99** | **85.45** | **49.80** | **83.22** |
| PCA | 75.09 | 71.10 | 63.51 | 76.60 | 67.36 | 74.77 |
| KPCA | **59.03** | **80.06** | **51.69** | **82.46** | **54.14** | **81.66** |
| HE | 59.07 | 78.95 | 64.12 | 76.92 | 62.44 | 77.59 |
| KHE | **57.80** | **80.72** | **54.42** | **80.72** | **55.53** | **80.72** |

### 5.4 Analyzing parameter sensitivity of the kernel

In this section, we analyze the cerebellar kernel’s key parameters, which determine the number of elements preserved in the projected vector for the explicit feature mapping Ψ_ceb_(*x*). The nonlinearity arises entirely from Top-k truncation, which reduces inter-sample similarity by removing low-activation components that do not contribute to discrimination. As shown in Figure 2, varying *s* on ImageNet-200 reveals a trade-off: very small *s* over-separates ID and OOD samples, while very large s fails to sufficiently reduce correlations. Optimal performance occurs at intermediate *s*, balancing nonlinearity with preservation of semantic information.

**Figure 2:**
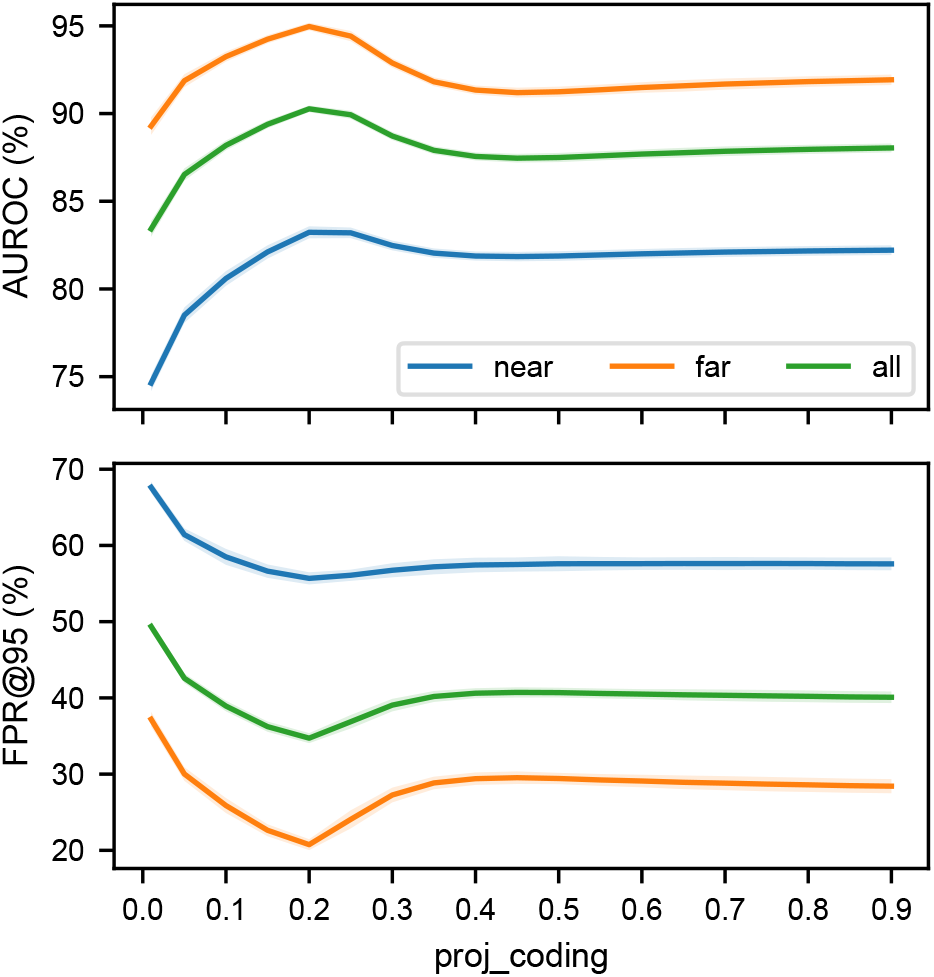
Effect of retention ratios *s* on OOD detection for ImageNet-200.

When designing the cerebellar kernel, considering differences between biological features and those in deep learning models, we developed the Abs-Topk variant. This design leverages representative information from positive and negative extremes, potentially better suited to machine learning features. We compare Abs-Topk with Pos-Topk on ImageNet-200 (Table 4) and find that Abs-Topk achieves better AUROC across datasets, supporting the effectiveness of our design.

**Table 4:** AUROC comparison of Abs-Topk and Pos-Topk settings on ImageNet-200.

| Setting | Near-OOD | Far-OOD | ALL |
| --- | --- | --- | --- |
| Pos-Topk | 82.88 | 94.74 | 89.99 |
| Abs-Topk | **83.23** | **94.96** | **90.27** |

### 5.5 Explicit mapping vs. closed-form computation

A primary goal of the cerebellar kernel is to avoid the high computational cost and variability introduced by explicitly computing random-projection features at large expansion dimensions. The closed-form kernel offers a deterministic alternative, corresponding to the theoretically optimal limit as the expansion dimension approaches infinity. In this section, we compare the closed-form CKE with an explicit-mapping variant that replaces the cerebellar kernel with its explicit feature mapping, highlighting the performance and computational advantages of closed-form computation.

As shown in Table 5, performance for explicit mapping improves monotonically with the expansion ratio, but marginal gains diminish rapidly—doubling the expansion yields only minor improvements. At sufficiently large expansion, explicit mapping approaches the performance of the closed-form model, yet remains slightly lower.

**Table 5:** Performance comparison between closed-form computation and explicit mapping across different expansion factors.

| Expansion | FPR@95 | AUROC |
| --- | --- | --- |
| 5× | 38.91 | 88.59 |
| 10× | 36.31 | 89.89 |
| 20× | 35.57 | 89.95 |
| 40× | 34.95 | 90.10 |
| Closed-form | **34.70** | **90.27** |

From the computational perspective, the cerebellar kernel reduces the per-pair complexity from *O*(*MN*) in explicit computation to *O*(*N*), where *M* is the expansion dimension and *N* is the original feature dimension. In practice, the end-to-end speedup may be smaller than the theoretical factor *M* due to other pipeline components and hardware utilization. As summarized in Table 6, in the forward pass, closed-form CKE is *13*.*7* × faster than the explicit mapping model with 20× expansion, and *24*.*6* × faster with 40× expansion.

**Table 6:** Comparison of forward-pass speed: explicit mapping vs. closed-form computation.

| Expansion | Explicit | Closed-form | Speedup |
| --- | --- | --- | --- |
| 20× | 25.53 | 1.86 | 13.7 |
| 40× | 46.29 | 1.88 | **24.6** |

## 6 Conclusion

This paper builds on the cerebellum-inspired structure of “random projection + Topk gating” and derives a practical, engineering-friendly kernel function. By replacing explicit high-dimensional mapping with a closed-form cerebellar kernel, we incorporate the geometric properties of cerebellar space into the post-processing stage of OOD detection without incurring high computational costs or randomness-induced instability. The resulting CKE achieves stable and substantial gains across datasets and settings. Moreover, the cerebellar kernel improves the performance of diverse methods, indicating that it is not a method-specific optimization but rather provides a more general geometric separation by reshaping feature similarity, thereby easing OOD detection.

## 7 Impact Statement

This work introduces a closed-form, cerebellum-inspired kernel that embeds the geometric properties of cerebellar space into OOD detection, eliminating the computational cost and instability of explicit high-dimensional mapping. The resulting CKE delivers consistent gains across benchmarks and improves diverse methods, demonstrating that it is a general mechanism for feature-space separation rather than a method-specific tweak. By efficiently reshaping feature similarity to enhance ID/OOD separability, this approach provides a scalable tool for robust pattern separation in OOD detection and related tasks.

## A Additional Derivations

### A.1 Top-k threshold and quantile approximation

In the explicit mapping Ψ_ceb_(*x*) = Topk(*Wx*), Top-k gating keeps the *k* = *sM* largest responses among 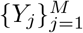. For analysis, we approximate this empirical Top-k threshold by a population quantile threshold *τ* (*x*) satisfying Pr(*Y* ≥ *τ*) = *s*. Under mild regularity conditions, the empirical Top-k threshold converges to the corresponding population quantile as *M* → ∞, justifying the quantile-threshold approximation used in Section 3.

### A.2 Integral forms of truncated moments

For (*Z, Z*′) ∼ 𝒩(0, Σ_*ρ*_) with correlation *ρ* ∈ (−1, 1), define *I* = **1**{*Z* ≥ *a*} and *I*′ = **1**{*Z*′ ≥ *b*}. Let

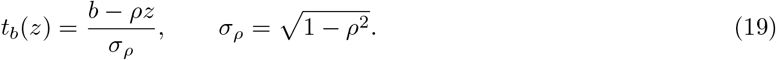

Then the two-dimensional truncated moments admit one-dimensional integral representations:

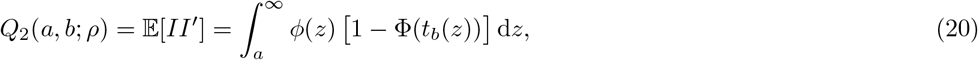

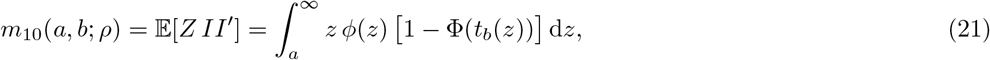

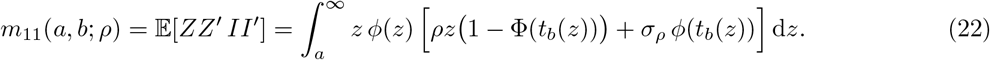

In the main text we use standardized gating where the threshold depends only on the retention ratio, so typically *a* = *b*.

### A.3 Abs-Topk moment identities

For Abs-Topk with standardized magnitude gating, let *I* = **1**{|*Z*| ≥ *a*} where *a* = Φ^−1^(1 − *s*/2) so that Pr(|*Z*| ≥ *a*) = *s*. By symmetry,

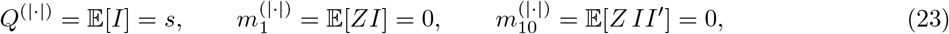

and

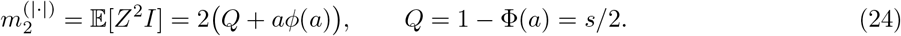

The two-dimensional truncated moments under |*Z*| gating can be obtained by combining one-sided moments at *ρ* and −*ρ*:

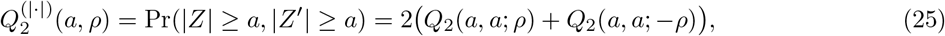

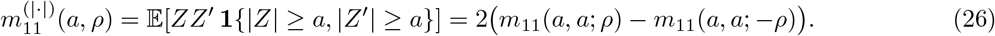

This yields the Abs-Topk ingredients 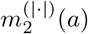 and 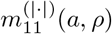 used in Section 4.1. An alternative Abs-Topk definition based on raw magnitude |*Y*| leads to thresholds that vary with the scale of *Y* and is therefore less convenient for efficient tabulation.

## B Details of KPCA and KHE Baselines

This appendix provides the implementation details omitted in Section 4.3 for the subspace-based baseline (KPCA) and the memory-retrieval baseline (KHE). Both methods operate on penultimate-layer features extracted from the pretrained classifier; we denote the feature of an input sample by *x* ∈ ℝ^*d*^. In our kernelized versions, we replace the Euclidean inner product / cosine similarity by the cerebellar kernel *k*(*x, x*′) (Section 4.1), typically using 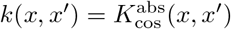.

### B.1 Kernel PCA residual OOD detector (KPCA)

#### Training (kernel principal subspace)

Let 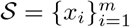 be a subset of correctly classified ID training features (we subsample to control memory and runtime). Define the kernel matrix

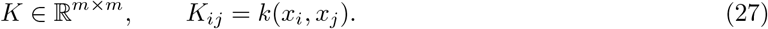

We optionally center the kernel in RKHS using the centering matrix 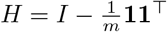 :

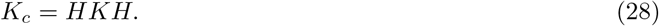

Let 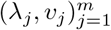 be the eigenpairs of *K*_*c*_ with *λ*_1_ ≥ *λ*_2_ ≥ · · · ≥ 0. Keeping the top *r* components, we form the normalized KPCA coefficients

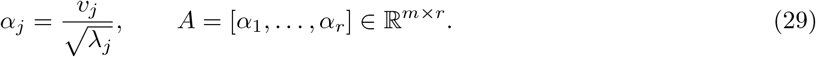

#### Scoring (reconstruction residual in kernel space)

For a test feature *x*, define the kernel vector to the subset

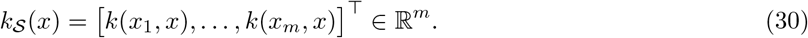

If centering is enabled, we use the centered test kernel vector

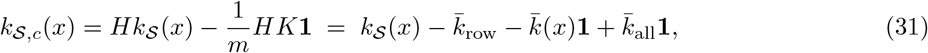

where 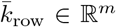 is the vector of row means of *K*, 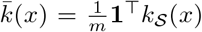, and 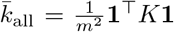. The KPCA embedding (principal-subspace coordinates) is then

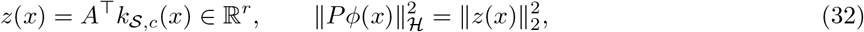

where *ϕ*(·) denotes the implicit RKHS feature map and *P* denotes the orthogonal projector onto the KPCA principal subspace. The centered self-kernel is

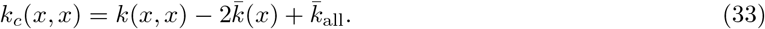

Finally, the KPCA reconstruction residual in RKHS is

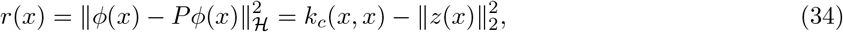

and we use the OOD confidence score *S*_KPCA_(*x*) = −*r*(*x*) (higher is more ID-like).

#### Class-specific KPCA for efficiency

To reduce the *O*(*m*) cost per sample, we also fit class-specific KPCA models 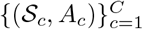 and score each test sample using the predicted-class subspace, i.e., *S*_KPCA_(*x*) = −*r*_*ĉ*(*x*)_(*x*).

### B.2 Kernel Hierarchical Energy (KHE)

#### Hopfield Energy (HE) as store-then-compare

HE [29] detects OOD samples by comparing a test feature against memorized ID *patterns*. Concretely, it stores a set of ID features (often organized per class). For a test feature *x* with predicted class *ĉ*(*x*), HE aggregates the similarities to the stored patterns of that class via a smooth max (log-sum-exp) with inverse temperature *β >* 0:

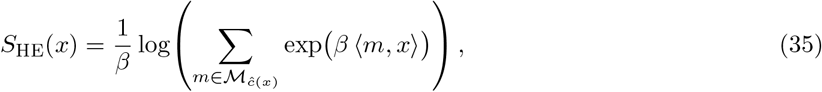

so that larger scores indicate higher similarity to memorized ID patterns (more ID-like), while smaller scores suggest OOD. Compared with Energy that pools over classifier logits, HE performs pooling over *pattern similarities* in feature space, which can be interpreted through the lens of modern Hopfield associative memory retrieval.

#### Kernelized variant (KHE)

KHE replaces the inner product ⟨*m, x*⟩ in HE by a kernel similarity *k*(*m, x*) (in our case, the cerebellar kernel from Section 4.1), yielding a memory-retrieval score that operates in the induced RKHS while keeping the same log-sum-exp aggregation. Let 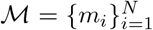 be the memory bank constructed from correctly classified ID training samples (stored either globally or per class). Given a test feature *x*, we compute similarities *k*(*m*_*i*_, *x*) and define the kernel-energy score

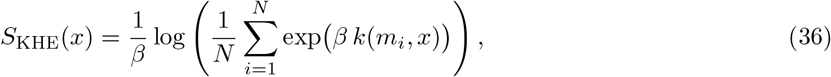

where *β >* 0 is an inverse-temperature parameter. We also evaluate a class-conditional variant by restricting ℳ to the predicted class *ĉ*(*x*), which reduces computation and can improve robustness when classes are well separated.

## C Full ImageNet Tables

**Table 7:** Full results on the ImageNet-200 Standard OOD benchmark.

| Method | SSB-hard |  | NINCO |  | Near-OOD |  | iNaturalist |  | Textures |  | OpenImage-O |  | Far-OOD |  | AL |
| --- | --- | --- | --- | --- | --- | --- | --- | --- | --- | --- | --- | --- | --- | --- | --- |
|  | FPR@95 | AUROC | FPR@95 | AUROC | FPR@95 | AUROC | FPR@95 | AUROC | FPR@95 | AUROC | FPR@95 | AUROC | FPR@95 | AUROC | FPR@95 |
| OpenMax | 72.37 | 77.53 | 54.59 | 83.01 | 63.48 | 80.27 | 24.53 | 92.32 | 36.80 | 90.21 | 38.03 | 88.07 | 33.12 | 90.20 | 45.26 |
| ODIN | 73.51 | 77.19 | 60.00 | 83.34 | 66.76 | 80.27 | 22.39 | 94.37 | 42.99 | 90.65 | 37.30 | 90.11 | 34.23 | 91.71 | 47.24 |
| MDS | 83.65 | 58.38 | 74.57 | 65.48 | 79.11 | 61.93 | 58.53 | 75.03 | 58.16 | 79.25 | 68.29 | 69.87 | 61.66 | 74.72 | 68.64 |
| MDSEns | 92.13 | 50.46 | 91.36 | 58.18 | 91.75 | 54.32 | 83.37 | 62.16 | 72.27 | 80.70 | 87.26 | 64.96 | 80.96 | 69.27 | 85.28 |
| RMDS | **65.91** | 80.20 | **42.13** | 84.94 | **54.02** | 82.57 | 24.70 | 90.64 | 37.80 | 86.77 | 34.85 | 86.77 | 32.45 | 88.06 | 41.08 |
| Gram | 85.68 | 65.95 | 87.13 | 69.40 | 86.40 | 67.67 | 85.54 | 65.30 | 80.87 | 80.53 | 86.66 | 67.72 | 84.36 | 71.19 | 85.18 |
| EBO | 69.77 | 79.83 | 50.70 | 85.17 | 60.24 | 82.50 | 26.41 | 92.55 | 41.43 | 90.79 | 36.74 | 89.23 | 34.86 | 90.86 | 45.01 |
| ReAct | 71.51 | 78.97 | 53.47 | 84.76 | 62.49 | 81.87 | 22.97 | 93.65 | 29.67 | 92.86 | 32.86 | 90.40 | 28.50 | 92.31 | 42.10 |
| VIM | 71.28 | 74.04 | 47.10 | 83.32 | 59.19 | 78.68 | 27.34 | 90.96 | 20.39 | 94.61 | 33.86 | 88.20 | 27.20 | 91.26 | 40.00 |
| KNN | 73.71 | 77.03 | 46.64 | 86.10 | 60.18 | 81.57 | 24.46 | 93.99 | 24.45 | 95.29 | 32.90 | 90.19 | 27.27 | 93.16 | 40.43 |
| ASH | 72.14 | 79.52 | 57.63 | 85.24 | 64.89 | 82.38 | 22.49 | 95.10 | 25.65 | 94.77 | 33.72 | 91.82 | 27.29 | 93.90 | 42.33 |
| SHE | 72.64 | 78.30 | 60.96 | 82.07 | 66.80 | 80.18 | 34.38 | 91.43 | 45.58 | 90.51 | 46.54 | 87.49 | 42.17 | 89.81 | 52.02 |
| GEN | 66.79 | **80.75** | 43.61 | 86.60 | 55.20 | **83.68** | 22.03 | 93.70 | 42.01 | 90.25 | 32.25 | 90.13 | 32.10 | 89.81 | 41.34 |
| CKE | 68.62 | 79.46 | 42.74 | **86.99** | 55.68 | 83.23 | **16.08** | **96.12** | **19.26** | **96.28** | **26.89** | **92.48** | **20.75** | **94.96** | **34.70** |

**Table 8:** Full results on the ImageNet-1k Standard OOD benchmark.

| Method | SSB-hard |  | NINCO |  | Near-OOD |  | iNaturalist |  | Textures |  | OpenImage-O |  | Far-OOD |  | AL |
| --- | --- | --- | --- | --- | --- | --- | --- | --- | --- | --- | --- | --- | --- | --- | --- |
|  | FPR@95 | AUROC | FPR@95 | AUROC | FPR@95 | AUROC | FPR@95 | AUROC | FPR@95 | AUROC | FPR@95 | AUROC | FPR@95 | AUROC | FPR@95 |
| OpenMax | 77.33 | 71.37 | 60.81 | 78.17 | 69.07 | 74.77 | 25.29 | 92.05 | 40.26 | 88.10 | 37.39 | 87.62 | 34.31 | 89.26 | 48.21 |
| ODIN | 76.83 | 71.74 | 68.16 | 77.77 | 72.50 | 74.75 | 35.98 | 91.17 | 49.24 | 89.00 | 46.67 | 88.23 | 43.96 | 89.47 | 55.38 |
| MDS | 92.10 | 48.50 | 78.80 | 62.38 | 85.45 | 55.44 | 73.81 | 63.67 | 42.79 | 89.80 | 72.15 | 69.27 | 62.92 | 74.25 | 71.93 |
| MDSEns | 95.19 | 43.92 | 91.86 | 55.41 | 93.52 | 49.67 | 84.23 | 61.82 | 73.31 | 79.94 | 90.77 | 60.80 | 82.77 | 67.52 | 87.07 |
| RMDS | 77.88 | 71.77 | 52.20 | 82.22 | 65.04 | 76.99 | 33.67 | 87.24 | 48.80 | 86.08 | 40.27 | 85.84 | 40.91 | 86.38 | 50.56 |
| Gram | 89.39 | 57.39 | 83.87 | 66.01 | 86.63 | 61.70 | 67.89 | 76.67 | 58.80 | 88.02 | 75.39 | 74.43 | 67.36 | 79.71 | 75.07 |
| EBO | 76.54 | 72.08 | 60.58 | 79.70 | 68.56 | 75.89 | 31.30 | 90.63 | 45.77 | 88.70 | 38.09 | 89.06 | 38.39 | 89.47 | 50.46 |
| ReAct | 77.55 | 73.03 | 55.82 | 81.73 | 66.69 | 77.38 | 16.72 | 96.34 | 29.64 | 92.79 | 32.58 | 91.87 | 26.31 | 93.67 | 42.46 |
| VIM | 80.41 | 65.54 | 62.29 | 78.63 | 71.35 | 72.08 | 30.68 | 89.56 | 10.51 | **97.97** | 32.82 | 90.50 | 24.67 | 92.68 | 43.34 |
| KNN | 83.36 | 62.57 | 58.39 | 79.64 | 70.87 | 71.10 | 40.80 | 86.41 | 17.31 | 97.09 | 44.27 | 87.04 | 34.13 | 90.18 | 48.83 |
| ASH | **73.66** | 72.89 | 52.97 | 83.45 | 63.32 | 78.17 | 14.04 | 97.07 | 15.26 | 96.90 | 29.15 | 93.26 | 19.49 | 95.74 | 37.02 |
| SHE | 76.30 | 71.08 | 69.72 | 76.49 | 73.01 | 73.78 | 34.06 | 92.65 | 35.27 | 93.60 | 55.02 | 86.52 | 41.45 | 90.92 | 54.07 |
| GEN | 75.73 | 72.01 | 54.90 | 81.70 | 65.32 | 76.85 | 26.10 | 92.44 | 46.22 | 87.59 | 34.50 | 89.26 | 35.61 | 89.76 | 47.49 |
| CKE | 76.32 | **76.12** | **48.87** | **84.74** | **62.59** | **80.43** | **6.27** | **98.52** | **9.47** | 97.89 | **28.05** | **93.49** | **14.59** | **96.63** | **33.79** |

## D Full FSOOD Tables

**Table 9:** Full results on the ImageNet-200 FSOOD benchmark.

|  | SSB-hard |  | NINCO |  | Near-OOD |  | iNaturalist |  | Textures |  | OpenImage-O |  | Far-OOD |  | ALL |
| --- | --- | --- | --- | --- | --- | --- | --- | --- | --- | --- | --- | --- | --- | --- | --- |
| Method | FPR@95 | AUROC | FPR@95 | AUROC | FPR@95 | AUROC | FPR@95 | AUROC | FPR@95 | AUROC | FPR@95 | AUROC | FPR@95 | AUROC | FPR@95 AU |
| OpenMax | 91.55 | 47.64 | 85.00 | 54.15 | 88.27 | 50.89 | 68.19 | 72.44 | 76.72 | 69.12 | 77.39 | 62.31 | 74.10 | 67.96 | 79.77 |
| ODIN | 91.71 | 44.31 | 86.66 | 52.36 | 89.18 | 48.33 | 65.50 | 70.19 | 79.13 | 67.10 | 76.03 | 61.48 | 73.55 | 66.25 | 79.81 |
| MDS | 93.90 | 48.59 | 88.52 | 56.65 | 91.21 | 52.62 | 74.46 | 68.25 | 74.10 | 73.84 | 83.68 | 61.90 | 77.41 | 68.00 | 82.93 |
| MDSEns | 96.28 | 34.22 | 95.83 | 41.58 | 96.06 | 37.90 | 91.39 | 43.63 | 84.83 | 67.54 | 93.52 | 48.38 | 89.91 | 53.18 | 92.37 |
| RMDS | 89.44 | 56.24 | **79.23** | 60.95 | **84.33** | 58.59 | 65.62 | 71.71 | 76.73 | 64.61 | 74.82 | 63.52 | 72.39 | 66.62 | 77.17 |
| Gram | **86.37** | **59.12** | 87.59 | 63.35 | 86.98 | **61.23** | 86.24 | 58.42 | 82.76 | 75.86 | 87.23 | 61.51 | 85.41 | 65.26 | 86.04 |
| EBO | 90.71 | 47.56 | 83.61 | 53.45 | 87.16 | 50.51 | 70.53 | 65.91 | 79.46 | 68.03 | 77.14 | 59.83 | 75.71 | 64.59 | 80.29 |
| ReAct | 91.22 | 47.25 | 84.61 | 53.84 | 87.91 | 50.55 | 67.52 | 69.45 | 72.82 | 71.45 | 74.81 | 62.30 | 71.72 | 67.73 | 78.20 |
| VIM | 91.61 | 45.34 | 82.35 | 57.09 | 86.98 | 51.22 | 68.15 | 71.34 | 58.50 | 82.54 | 74.54 | 65.70 | 67.06 | 73.19 | 75.03 |
| KNN | 91.73 | 44.05 | 81.23 | 54.51 | 86.48 | 49.28 | 69.10 | 71.53 | 69.06 | 81.88 | 74.43 | 62.12 | 70.86 | 71.84 | 77.11 |
| ASH | 90.29 | 50.96 | 84.21 | 58.51 | 87.25 | 54.74 | **63.16** | 77.96 | 65.99 | 79.39 | 71.69 | 69.09 | 66.95 | 75.48 | 75.07 |
| SHE | 91.16 | 52.82 | 86.49 | 56.64 | 88.82 | 54.73 | 71.48 | 72.20 | 78.98 | 74.27 | 79.50 | 64.95 | 76.65 | 70.47 | 81.52 |
| GEN | 89.59 | 48.33 | 80.12 | 54.85 | 84.85 | 51.59 | 66.47 | 68.94 | 79.30 | 66.58 | 73.96 | 60.87 | 73.24 | 65.46 | 77.89 |
| NAC | 92.75 | 45.42 | 88.83 | 53.80 | 90.79 | 49.61 | 72.57 | 65.83 | 69.08 | 74.41 | 79.55 | 60.79 | 73.73 | 67.01 | 80.56 |
| KAN | 94.67 | 58.37 | 87.35 | 61.10 | 91.01 | 59.74 | 64.89 | **84.13** | 68.75 | 83.30 | 78.84 | 70.40 | 70.83 | 79.28 | 78.90 |
| CKE | 90.73 | 52.09 | 80.41 | **63.56** | 85.57 | 57.83 | 84.83 | 59.28 | **57.70** | **88.48** | **71.02** | **75.23** | **61.21** | **82.85** | **70.95** |

**Table 10:** Full results on the ImageNet-1k FSOOD benchmark.

|  | SSB-hard |  | NINCO |  | Near-OOD |  | iNaturalist |  | Textures |  | OpenImage-O |  | Far-OOD |  | ALL |
| --- | --- | --- | --- | --- | --- | --- | --- | --- | --- | --- | --- | --- | --- | --- | --- |
| Method | FPR@95 | AUROC | FPR@95 | AUROC | FPR@95 | AUROC | FPR@95 | AUROC | FPR@95 | AUROC | FPR@95 | AUROC | FPR@95 | AUROC | FPR@95 AU |
| OpenMax | 86.76 | 53.79 | 76.41 | 60.28 | 81.59 | 57.03 | 49.98 | 80.30 | 62.28 | 73.54 | 60.05 | 71.88 | 57.44 | 75.24 | 67.10 |
| ODIN | 86.01 | 54.22 | 80.40 | 60.59 | 83.21 | 57.41 | 57.10 | 77.43 | 67.34 | 76.04 | 65.45 | 73.40 | 63.29 | 75.62 | 71.26 |
| MDS | 95.41 | 39.22 | 86.89 | 52.83 | 91.15 | 46.02 | 83.38 | 54.06 | 55.95 | 86.26 | 82.16 | 60.75 | 73.83 | 67.02 | 80.76 |
| MDSEns | 96.22 | 37.13 | 93.56 | 47.80 | 94.89 | 42.47 | 87.47 | 53.32 | 78.56 | 73.39 | 92.72 | 53.24 | 86.25 | 59.98 | 89.71 |
| RMDS | 87.09 | 56.61 | 70.28 | 67.50 | 78.69 | 62.06 | 55.68 | 73.48 | 67.83 | 74.25 | 61.30 | 72.13 | 61.61 | 73.29 | 68.44 |
| Gram | 90.87 | 51.93 | 86.24 | 60.63 | 88.55 | 56.28 | 72.69 | 71.36 | 64.65 | 84.83 | 79.05 | 69.40 | 72.13 | 75.20 | 78.70 |
| EBO | 86.19 | 52.93 | 76.15 | 60.28 | 81.17 | 56.61 | 55.18 | 74.01 | 66.05 | 73.89 | 60.55 | 72.22 | 60.59 | 73.37 | 68.82 |
| ReAct | 86.90 | 55.34 | 72.97 | 64.51 | 79.94 | 59.93 | 40.64 | 87.93 | 53.47 | 81.08 | 55.89 | 79.34 | 50.00 | 82.78 | 61.97 |
| VIM | 88.85 | 45.88 | 77.85 | 59.12 | 83.35 | 52.50 | 55.59 | 72.22 | 33.64 | 93.09 | 57.23 | 75.01 | 48.82 | 80.10 | 62.63 |
| KNN | 90.50 | 43.78 | 74.77 | 59.86 | 82.64 | 51.82 | 62.46 | 67.79 | 42.59 | 90.29 | 65.03 | 69.98 | 56.69 | 76.02 | 67.07 |
| ASH | **84.49** | 54.66 | 70.74 | 66.38 | **77.61** | 60.52 | 36.82 | 89.23 | 38.34 | 89.53 | 52.33 | 81.47 | 42.50 | 86.75 | 56.54 |
| SHE | 85.10 | 58.15 | 80.47 | 64.27 | 82.78 | 61.21 | 50.97 | 84.71 | 52.17 | 87.48 | 69.38 | 76.92 | 57.50 | 83.04 | 67.62 |
| GEN | 85.72 | 52.95 | 72.38 | 62.73 | 79.05 | 57.84 | 50.61 | 78.47 | 66.40 | 71.82 | 57.67 | 72.62 | 58.23 | 74.31 | 66.56 |
| NAC | 89.54 | 52.48 | 76.53 | 66.49 | 83.03 | 59.48 | 36.88 | 88.92 | **29.19** | 92.77 | 58.48 | 80.76 | 41.52 | 87.48 | 58.12 |
| KAN | 89.64 | 55.88 | 74.57 | 69.55 | 82.11 | 62.71 | 42.67 | 91.55 | 37.79 | 93.45 | 67.66 | 82.15 | 49.37 | 89.05 | 62.47 |
| CKE | 86.41 | **61.08** | **69.04** | **71.00** | 77.72 | **66.04** | **18.23** | **95.36** | **24.95** | **94.35** | **51.58** | **85.20** | **31.59** | **91.64** | **50.04** |

## E Full CIFAR-100 Table

**Table 11:** Full results on the CIFAR-100 benchmark.

| Method | CIFAR-10 |  | TIN |  | Near-OOD |  | MNIST |  | SVHN |  | Textures |  | Places365 |  | Far-OOD |  | ALL |  |
| --- | --- | --- | --- | --- | --- | --- | --- | --- | --- | --- | --- | --- | --- | --- | --- | --- | --- | --- |
|  | FPR@95 | AUROC | FPR@95 | AUROC | FPR@95 | AUROC | FPR@95 | AUROC | FPR@95 | AUROC | FPR@95 | AUROC | FPR@95 | AUROC | FPR@95 | AUROC | FPR@95 | AUROC |
| OpenMax | 60.17 | 74.38 | 52.99 | 78.44 | 56.58 | 76.41 | 53.82 | 76.01 | 53.20 | 82.07 | 56.12 | 80.56 | 54.85 | 79.29 | 54.50 | 79.48 | 55.19 | 78.46 |
| ODIN | 60.64 | 78.18 | 55.19 | 81.63 | 57.91 | 79.90 | 45.94 | 83.79 | 67.41 | 74.54 | 62.37 | 79.33 | 59.71 | 79.45 | 58.86 | 79.28 | 58.54 | 79.49 |
| MDS | 88.00 | 55.87 | 79.05 | 61.50 | 83.53 | 58.69 | 71.72 | 67.47 | 67.21 | 70.68 | 70.49 | 76.26 | 79.61 | 63.15 | 72.26 | 69.39 | 76.01 | 65.82 |
| MDSEns | 95.94 | 43.85 | 95.82 | 48.78 | 95.88 | 46.31 | **2.83** | **98.21** | 82.57 | 53.76 | 84.94 | 69.75 | 96.61 | 42.27 | 66.74 | 66.00 | 76.45 | 59.44 |
| RMDS | 61.37 | 77.75 | 49.56 | 82.55 | 55.46 | 80.15 | 52.05 | 79.74 | 51.65 | 84.89 | 53.99 | 83.65 | 53.57 | 83.40 | 52.81 | 82.92 | 53.70 | 82.00 |
| Gram | 92.71 | 49.41 | 91.85 | 53.91 | 92.28 | 51.66 | 53.53 | 80.71 | **20.06** | **95.55** | 89.51 | 70.79 | 94.67 | 46.38 | 64.44 | 73.36 | 73.72 | 66.12 |
| EBO | 59.21 | 79.05 | 52.03 | 82.76 | 55.62 | 80.91 | 52.62 | 79.18 | 53.62 | 82.03 | 62.35 | 78.35 | 57.75 | 79.52 | 56.59 | 79.77 | 56.26 | 80.15 |
| ReAct | 61.30 | 78.65 | 51.47 | 82.88 | 56.39 | 80.77 | 56.04 | 78.37 | 50.41 | 83.01 | 55.04 | 80.15 | 55.30 | 80.03 | 54.20 | 80.39 | 54.93 | 80.52 |
| VIM | 70.59 | 72.21 | 54.66 | 77.76 | 62.63 | 74.98 | 48.32 | 81.89 | 46.22 | 83.14 | 46.86 | 85.91 | 61.57 | 75.85 | 50.74 | 81.70 | 54.70 | 79.46 |
| KNN | 72.80 | 77.02 | 49.65 | **83.34** | 61.22 | 80.18 | 48.58 | 82.36 | 51.75 | 84.15 | 53.56 | 83.66 | 60.70 | 79.43 | 53.65 | 82.40 | 56.17 | 81.66 |
| ASH | 68.06 | 76.48 | 63.35 | 79.92 | 65.71 | 78.20 | 66.58 | 77.23 | 46.00 | 85.60 | 61.27 | 80.72 | 62.95 | 78.76 | 59.20 | 80.58 | 61.37 | 79.79 |
| SHE | 60.41 | 78.15 | 57.74 | 79.74 | 59.07 | 78.95 | 58.78 | 76.76 | 59.15 | 80.97 | 73.29 | 73.64 | 65.24 | 76.30 | 64.12 | 76.92 | 62.44 | 77.59 |
| GEN | **58.87** | **79.38** | 49.98 | 83.25 | **54.42** | **81.31** | 53.92 | 78.29 | 55.45 | 81.41 | 61.23 | 78.74 | 56.25 | 80.28 | 56.71 | 79.68 | 55.95 | 80.23 |
| CKE | 61.72 | 75.64 | **49.14** | 81.88 | 55.43 | 78.76 | 52.06 | 76.84 | 40.01 | 90.93 | **43.81** | **89.75** | **52.08** | **84.27** | **46.99** | **85.45** | **49.8** | **83.22** |

